# Antibody affinity engineering using antibody repertoire data and machine learning

**DOI:** 10.1101/2025.01.10.632313

**Authors:** Lena Erlach, Simon Friedensohn, Daniel Neumeier, Derek M. Mason, Sai T. Reddy

**Affiliations:** Department of Biosystems Science and Engineering, ETH Zürich, 4056 Basel, Switzerland; Botnar Institute of Immune Engineering, 4056 Basel, Switzerland

## Abstract

Advanced antibody discovery and engineering workflows take advantage of the combination of high-throughput screening, deep sequencing and machine learning (ML). Most high-throughput methods, however, lack the resolution to provide absolute affinity values of antibody-antigen interactions, limiting their utility for precise engineering of binding kinetics. In this study, we utilize antibody repertoire data, affinity characterization and ML for antibody affinity engineering. Leveraging natural antibody sequence information from repertoires of immunized mice, we identified and experimentally measured affinities for 35 antigen-specific variants. Supervised ML models trained on these sequences achieved remarkable accuracy in predicting affinity, despite the limited dataset size. We utilized the trained ML model to *in silico*-design eight synthetic antibody variants, of which seven exhibited the desired affinities. Our study illustrates the potential of this streamlined and efficient approach for precise engineering of the affinity of antibodies while reducing extensive experimental screening.

## Introduction

*In vivo* approaches relying on animal immunization with target antigens are widely used for antibody discovery for therapeutic, diagnostic and research purposes (Lu et al. 2020). Antibodies derived from B cells of immunized hosts are remarkably diverse and can exhibit high affinity and specificity, even against complex antigens (Finney et al. 2018). *In vivo-*derived antibodies are selected through highly regulated B cell immune processes, such as clonal selection, expansion and somatic hypermutation (SHM) (Mesin et al. 2016), and therefore often possess favorable drug developability properties when compared to antibodies derived from *in vitro* methods (e.g., phage or yeast display of recombinant libraries) (Prabakaran et al. 2021; Jain et al. 2017). Due to the unique advantage in retaining native heavy and light chain pairing, hybridoma screening or single B cell cloning are commonly used methods to isolate such antibodies (Kellermann & Green 2002; Laustsen et al. 2021; Prabakaran et al. 2021). However, these methods are constrained by experimental screening to ∼100-1,000 B cells from a single immunized mouse.

Advances in deep sequencing have revolutionized the study of antibody repertoires, enabling the identification of millions of antibody variants per individual and revealing the vast, untapped diversity within these repertoires. Sequencing the antibody variable regions of heavy (VH) and light (VL) chains provides detailed insights into clonal selection, expansion, SHM, and antigen-binding regions (paratopes) such as the complementarity-determining region 3 (CDR3) (Reddy et al. 2010; Greiff et al. 2020). Bulk sequencing offers high depth profiling of repertoire diversity, while single-cell approaches provide native pairing of the heavy and light chains, albeit at much lower throughput. These technologies have led to the discovery of broadly neutralizing antibodies (Kong et al. 2019; Robbiani et al. 2020) and optimization of antibody affinity and stability (Hsiao et al. 2019). Despite these advances, challenges remain including limited throughput of single-cell methods and the need for robust annotation and labelling of the antibody variants. To address this limitation, high-throughput methods to screen for specificity of antibody sequences from natural repertoires have been developed (Bowers et al. 2011; Parola et al. 2019; Teixeira et al. 2022).

High-throughput screening and deep sequencing have supported the emerging field of computational antibody engineering, most notably the use of machine learning (ML) to decipher sequence-function relationships of antibodies (Pertseva et al. 2021). A common workflow starts with screening antibody libraries for binding and non-binding to a target antigen, followed by deep sequencing to generate sequence information with binary labels (binding or non-binding to target antigen). Antibody protein sequences can then be encoded (e.g., one-hot encoding) and their binding or non-binding labels serve as inputs to train ML classifiers. Such supervised ML models can then be used to predict whether an antibody is binding or non-binding based on the antibody sequence, thus allowing *in silico* screening of a much larger antibody sequence space. Supervised ML approaches encompass a wide range of architectures, from simple models such as linear regression or support vector machines to larger and more complex models like ensemble models or neural networks. Several studies have illustrated the success of this workflow in antibody engineering such as improving the affinity or developability parameters of a therapeutic antibody (Mason et al. 2021; Liu et al. 2020), reducing non- and poly-specific variants of antibodies (Saksena et al. 2022; Makowski et al. 2022; Harvey et al. 2022) or evaluating the impact of noise and unlabeled data for antibody specificity predictions (Minot & Reddy 2024). In addition to supervised ML, recent advances in natural language processing (NLP) have introduced large language models (Rives et al. 4 2021; Madani et al. 2020; Brandes et al. 2022) to the domain of protein and antibody engineering. Protein language models (PLMs) are pre-trained on the extensive protein sequence databases in an unsupervised manner enabling them to capture implicit features of evolutionary fitness that are hypothesized to correlate with functional protein fitness. Based on this concept, PLMs have demonstrated notable success in guiding affinity maturation without explicit supervision, as shown by (Hie et al. 2023), who used these models to propose mutations that improved antibody affinity. Furthermore, PLMs can be augmented with structural information to enhance unsupervised affinity maturation (Shanker et al. 2024).

While methods based on binary labels from antibody display methods and unsupervised approaches with PLMs have proven successful in antibody engineering they rely on screening synthetic mutagenesis libraries of antibody variants. Here, we present a case study for antibody affinity engineering leveraging natural antibody repertoire datasets, which may have favorable developability properties, combined with ML models trained with small datasets of antibody-antigen affinity measurements (<50 variants). Importantly, in contrast to previously mentioned studies that use ML classification models to make binary predictions on antibody specificity (binding or non-binding), we use regression ML models capable of making predictions of continuous numerical values, such as an antibody’s binding affinity (K_D_ value). Starting from a known antibody sequence with specificity to a model antigen (hen egg lysozyme, HEL), we developed a computational workflow to identify other potential antigen-specific antibody variants from repertoire data of immunized mice. Recombinant antibody expression and subsequent experimental characterization confirmed antigen specificity of 35 selected antibody variants. Subsequently, ML regression models were trained on antibody-antigen affinity measurements and evaluated for their performance in predicting affinity based on antibody sequence. To further validate this approach to guide affinity engineering, we used the trained ML models to design novel, synthetic antibody variants. Accurate affinity predictions were achieved for seven out of eight synthetic variants. This study underscores the potential of combining antibody repertoire sequencing with ML for antibody affinity engineering.

## Results

### Sequence selection from antibody repertoires of murine plasma cell compartment

This study utilized antibody repertoire datasets previously described by Friedensohn et al. (Friedensohn et al. n.d.). Antibody repertoires from the bone marrow of 12 BALB/c mice, immunized with the model antigen HEL, were characterized using targeted (bulk) deep sequencing of antibody VH chains (Illumina Miseq, 2 x 300 bp). Mice were divided into four groups (n = 3 per group; see Supplementary Table 1), with each group receiving a different number of booster immunizations: zero, one, two, or three. The VH sequencing libraries were generated by extracting RNA and performing a two-step reverse transcription PCR (RT-PCR) that incorporated molecular barcoding to minimize errors and biases (Khan et al. 2016). The sequencing produced a total of 27,076 unique heavy chain complementary-determining region 3 (CDRH3) sequences and 83,479 unique VH sequences (full-length VDJ region).

As a starting sequence we selected a HEL-specific antibody VH variant (identified internally by experimental library screening), which we refer to as 3A-WT. Unlike single-cell sequencing, bulk repertoire data cannot preserve natural VH-VL pairings. Therefore 3A-WT was paired with a light chain from a previously identified anti-HEL antibody and demonstrated binding to HEL, which is in agreement with prior studies that have demonstrated that surrogate pairing of VL with VH-specific variants identified by deep sequencing can lead to antibody binding to target antigen (Reddy et al. 2010; Friedensohn et al. n.d.). Detailed information about the VH sequence and additional repertoire features of this variant are provided Supplementary Table 2. Beyond its confirmed binding specificity to HEL, the CDRH3 of 3A-WT was identified in more than one mouse repertoire dataset (Friedensohn et al. n.d.), suggesting that it represents a convergent clone, where there is a higher likelihood of similar sequences in these repertoires being antigen-specific (Greiff et al. 2015).

The first aim was to create a dataset of antibody variants from the antibody repertoire data that could be characterized experimentally and used to train and evaluate ML models for the *in silico* prediction of binding affinity to the target antigen. Consequently, the main focus of the sequence selection process was directed towards subsampling of the pooled antibody repertoire dataset from the 12 mice immunized with HEL consisting of 83,479 unique VH sequences (Supplementary Table 3). We aimed to express and characterize a single batch of 50 distinct variants.

The sequence selection process adhered to a well-defined workflow (Figure 2A). Initially, the repertoire dataset was filtered for VH CDR3 (CDRH3) amino acid (aa) sequences that were highly similar to the known 3A-WT variant’s CDRH3. To ensure a selection of potential antigen-binding variants, a conservative threshold of 80% aa similarity in Levenshtein distance (LD) was chosen based on the assumption that this is a common cutoff for assigning sequences to a clonal lineage (clonotype) of common antigen specificity (Greiff et al. 2015). In addition, affinity propagation (AP) clustering was employed (Frey & Dueck 2007). AP is a data-driven clustering method that does not require defining the number of clusters, and therefore allows for the intrinsic distribution of the dataset to guide the clustering. This method complements threshold-based filtering by leveraging global dataset characteristics to identify relevant sequence clusters beyond strict cutoffs. The vast majority of sequences (87%) identified by clonotyping and AP clustering were overlapping, suggesting that the 80% similarity threshold reflects the distribution of sequence cluster diversity in the repertoire dataset. A total of 26 CDRH3 sequences identified by both methods were selected for further analysis.

After filtering for CDRH3 sequences, the scope of the selection was extended to all VH sequences in the dataset that contained the selected CDRH3s, a total of 193 VH sequences. The selected VH sequences had between 1-14 SHMs, with a mean of 4.2. The overwhelming majority (94%) of sequences were aligned to IGHV1-15*01 and IGHJ3*01 germlines. Further details of the repertoire statistics are provided in Supplementary Table 4. Here, the goal was to identify antibody clonal variants that are likely to maintain antigen specificity, but possess varying affinities that will be experimentally measured. This dataset will then be used for training ML models that can predict antibody affinity based on antibody sequence. To this aim, k-medoids clustering was applied to the VH sequences and the cluster centroids were selected. K-medoids clustering is based on the principles of k-means clustering, but considers data points as cluster centers. This method was chosen based on the assumption that the cluster centers represent well distributed sample points of the full antigen-specific VH sequence space. The hyperparameter, k, is the number of clusters to which the data points are assigned. As the variant space of the dataset size was to be designed and selected to contain 50 variants (including the initial binding variant 3A-WT), k was set to 49. As anticipated, this sequence selection yielded a significant reduction in CDRH3 sequence diversity (Figure 2B) and a similarity network analysis (LD = 2) of the 193 VH sequences shows that most of the selected sequences (red nodes) are well distributed across the dataset and most clusters are represented among the selected variants (Figure 2C). The 50 selected antibody variants including the 3A-WT served as basis for generating the training dataset for modeling antibody-antigen binding by ML (Supplementary Table 5 and 6, Supplementary Figure 1).

**Figure 1.**
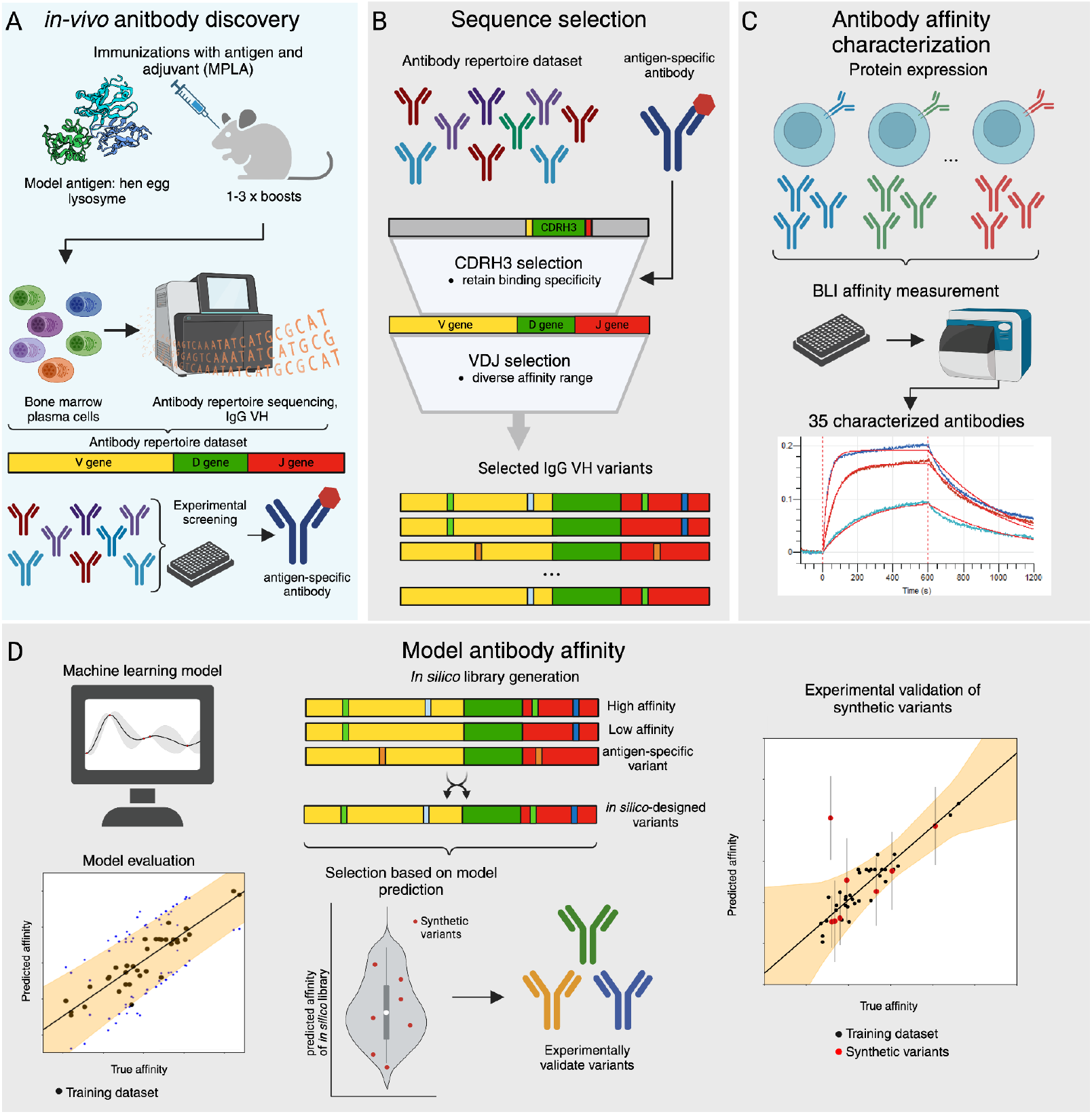
Overview figure of antibody affinity engineering using antibody repertoires and machine learning. (**A**) An antibody repertoire dataset was generated as described in Friedensohn et al. Bone marrow plasma cells were extracted and their antibody VH regions were sequenced after mouse immunization with the model antigen HEL. Through experimental screening, an antibody variant specific for HEL (3A-WT) was discovered internally. (**B**) Using this repertoire dataset, antibody VH sequences similar to the antigen-specific 3A-WT variant were selected. (**C**) The selected VH variants were expressed as soluble antibodies utilizing a mammalian display and expression platform. Binding affinity of the variants was determined by BLI (**D**) ML models were evaluated to predict binding affinity from antibody sequence in the training dataset. In order to experimentally validate the ML model predictions, synthetic antibody variants were designed based on the training dataset and model predictions. These synthetic, in silico-designed variants were expressed and characterized and confirmed high accuracy of the ML predictions.

**Figure 2.**
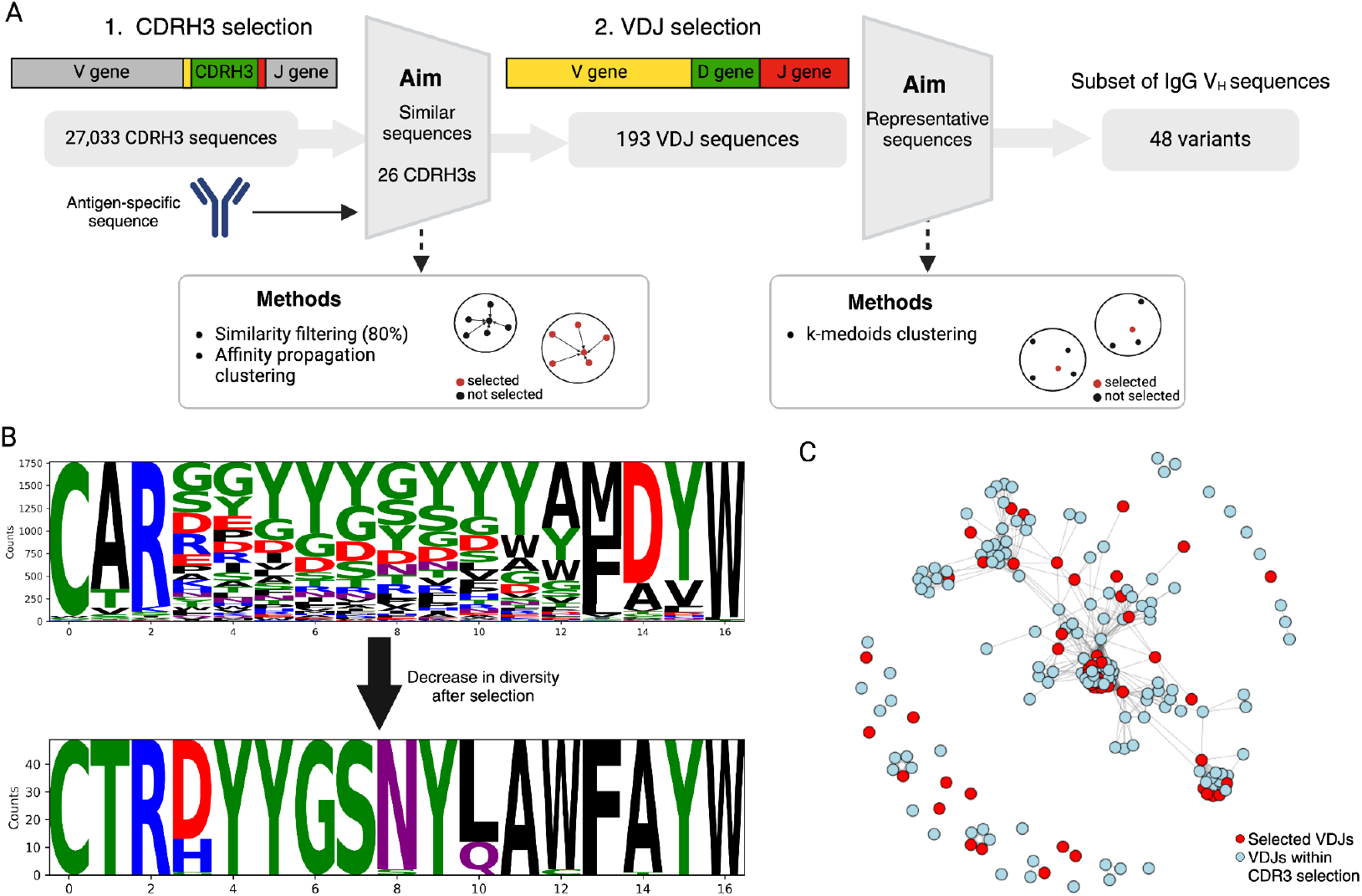
Selecting antibody VH variants. Detailed workflow of the VH sequence selection process is performed on two levels. (**A**) 1. Stringent CDRH3 selection is based on sequence similarity and affinity propagation (AP) clustering, which are combined to ensure antigen binding of the selected antibody variants. 2. Sequence selection based on VH (full VDJ region) uses k-medoids clustering to identify antibody variants that efficiently represent the sequence-space diversity and provide the potential to have varying affinities to target antigen. (**B**) Logoplots of the CDRH3 diversity before and after filtering. (**C**) Network plot depicting the sequence coverage of the selected VH sequences after filtering. Blue nodes show the full selection of VH sequences after the CDRH3 filtering and red nodes represent the selected sequences by VH filtering. Nodes are connected with edges when LD <= 2.

### Expression and experimental characterization of selected antibody variants

A previously developed mammalian cell antibody display and secretion system, Plug-n-Play (PnP) hybridomas (Pogson et al. 2016; Mason et al. 2018), was used as an expression platform for experimental characterization of the selected antibody variants (see Materials and Methods). Initial characterization of the antibody variants involved enzyme-linked immunosorbent assay (ELISA) screening of the supernatants to confirm antigen-specificity of the secreted antibodies. Notably, five variants (HC4, HC10, HC12, HC37 and HC45) did not show detectable binding to HEL antigen (could not be distinguished from the negative control) and were consequently excluded from further testing. The remaining 45 cell line variants were expanded in culture to generate sufficient soluble IgG.

Biolayer interferometry (BLI) was performed to measure binding kinetics and affinities [association rate (k_a_), dissociation rate (k_d_) and equilibrium dissociation constant (K_D_)] of the different antibody variants to HEL antigen (Supplementary Table 9, Supplementary Figure 3). To evaluate the reliability of the calculated kinetic coefficients, the coefficient of determination (R^2^ value) was computed. Among the 45 measured antibody variants, 10 were eliminated from the dataset, due to their measurements yielding R^2^ values below the defined threshold of 0.95. Low R^2^ values could be attributed to low expression levels or very low binding affinity towards the target antigen.

To visually examine the sequence-function relationship of the characterized antibody variants, both a network plot and a phylogenetic tree were generated (Figure 3). Nodes of the network plot represent a distinct antibody variant and edges (connections) correspond to variants with LD ≤ 2 (Figure 3A). Variants exhibiting higher connectivity tend to correspond to high binding affinities of K_D_ < 3 nM. The phylogenetic tree constructed through the neighbor-joining method (Saitou & Nei 1987) visualizes the mutational relation of the VH sequences and the corresponding affinities. The K_D_ values were categorized into bins of <1 nM, <3 nM, and >3 nM. One can observe that closely related sequences exhibit a tendency to share similar affinities. For example, the two variants with the lowest binding affinities, HC1 (K_D_ = 4.52 nM) and HC8 (K_D_ = 5.09) cluster together within the same sub-branch of the tree (Figure 3B). Furthermore, antibody sequences with K_D_ values above 1 nM and those with K_D_ values below 1 nM tend to segregate along the main branches of the tree. Notably, the number of SHMs in the VH sequence does not correlate with higher affinity.

**Figure 3.**
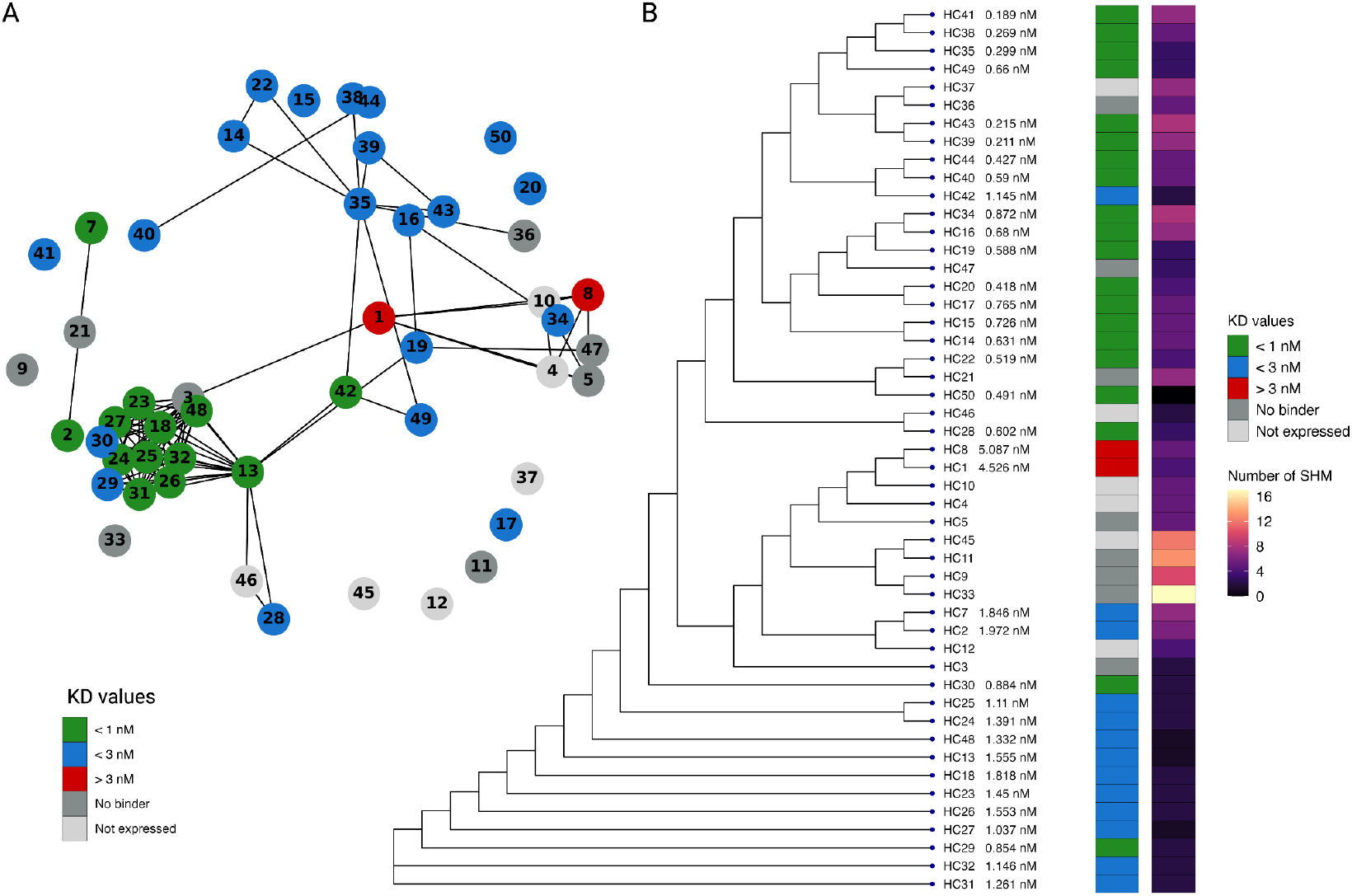
Characterization of antibody VH variants. (**A**) Network plot of the expressed and characterized antibody VH variants. Nodes are colored based on binned K_D_ values. (**B**) Phylogenetic tree representing the mutational relationship between the variants, K_D_ values and SHMs indicated by the color of the label.

### Machine learning enables accurate affinity prediction of antibody variants

We benchmarked five, linear and non-linear ML model architectures. The non-linear models included Gaussian Process (GP) models (Rasmussen et al. 2006) employing two distinct Gaussian kernels, specifically Matern kernel (GP_Matern) and Radial Basis Function (RBF) kernel (GP_RBF), Kernel-Ridge Regression (KRR) and Random Forest (RF). A Linear Regression (LR) model served as the linear baseline. To assess the performance of these models in a comprehensive manner, two cross-validation scenarios, (1) nested cross-validation (CV) and (2) leave-one-out CV (LOO-CV) were performed. Nested CV estimates the generalization error of the model including its (hyper)parameter search and therefore avoids overestimation of performance estimates. Our dataset however is small and LOO-CV is beneficial as it maximizes usage of the training dataset, and thus minimizes bias in the test set.

Within the nested CV experiments and relative to the other ML models, the GP_Matern and GP_RBF models performed better with R^2^ values of 0.7804 and 0.7589, respectively and mean-square error (MSE) of 0.0208 and 0.0225, respectively (Figure 4A and Supplementary Tables 10,11). LR performed the worst of the evaluated models and exhibited excessively large MSE estimates, affirming its inability to capture the non-linear relationships intrinsic to sequence-function associations. Therefore, LR was excluded from further detailed model comparisons.

**Figure 4.**
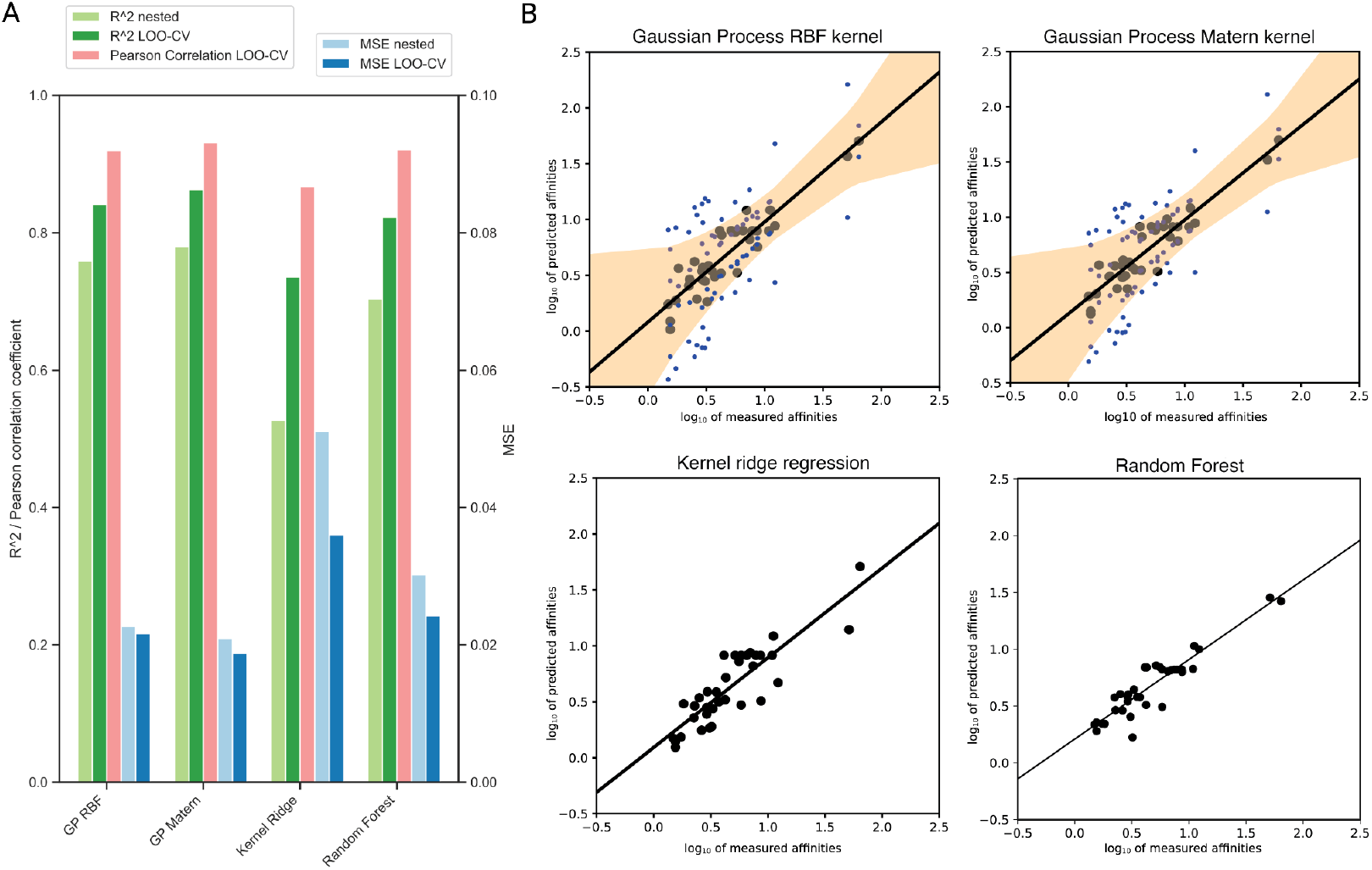
Comparative analysis of machine learning models and their performance on predicting antibody-antigen affinity. (**A**) Comparison of predictive performance of the benchmarked ML models in a nested cross-validation (CV) and leave-one-out CV (LOO-CV) framework (MSE, R^2^ and Pearson correlation coefficient). (**B**) Scatter plots depicting the correlation between true and predicted values for log_10_ antibody-antigen affinity from LOO-CV evaluations for the different regression models. The uncertainty estimate of the GP models is depicted as blue dots and the interpolated confidence interval (calculated as two times the standard deviation estimate of the GP models) is shown as the orange area around the correlation line.

In LOO-CV, the R^2^ and Pearson correlation coefficients were calculated using all predicted values of the training dataset, as these metrics cannot be assessed with single points. GP models again outperform RF and KRR with GP_Matern achieving the highest R^2^ value of 0.8625 and lowest MSE of 0.0188. In contrast to nested CV evaluations, the RF model in a LOO-CV scenario demonstrated regression metrics (R^2^=0.8224, MSE=0.0242) comparable to the GP models (Supplementary Table 10). ML models trained with randomized labels exhibited an inability to predict affinity from antibody sequences compared to models trained on the unaltered dataset (Supplementary Figure 4). This observation highlights that the chosen models effectively discern meaningful patterns for sequence-function prediction. Lastly, scatter plots of the predicted (LOO-CV) and true affinity values depict a robust correlation (Figure 4B). Notably, GPs introduce an additional layer of insight through their output of an uncertainty estimate for each prediction. This uncertainty estimate, which was calculated as two times the standard deviation estimate of the GP models, signifies the model’s confidence in its predictions. These findings underscore the potential of ML models for the prediction of antigen affinity from antibody sequences. While all models demonstrate good predictive capabilities, GPs exhibited enhanced performance, especially in small data scenarios.

### Design of affinity optimized antibody variants

In order to assess the predictive capabilities of the ML models on unseen antibody variants, we performed experimental validation with eight synthetic VH sequence variants that do not occur in the antibody repertoire dataset. These out-of-distribution variants were designed through an experimentally guided *in silico* mutagenesis approach and specifically selected to cover the observed affinity range. The mutational space was defined by five seed sequences, including the original antigen-specific variant (3A-WT) as well as the highest-(HC8, HC1) and lowest-affinity variants (HC41, HC39) from the training dataset. To minimize distant extrapolation and ensure reliable predictions, the synthetic variants were restricted to single or double point mutants of any of the seed sequences. Using the GP_Matern model trained on the training dataset, we selected eight variants that span the full range of predicted affinities (Figure 5A, Supplementary Table 12). Once again, these VH sequences were expressed as full-length antibodies using the PnP platform. The measured K_D_ values of the synthetic variants are comparable to the K_D_ values of the training dataset, confirming that they encompass the intended range of affinities (Figure 5B). We used these synthetic variants as the final test set for validating the predictive performance of the trained ML models (GP_RBF, GP_Matern, KRR, RF). R^2^, MSE and Pearson correlation coefficients of the K_D_ predictions of the synthetic variants were calculated (Supplementary Table 13). The predictive performance in this out-of-distribution scenario was reduced compared to training evaluation results (Figure 5C, Figure 4A, Supplementary Table 13). Among all ML models, the GP_Matern model demonstrated the highest predictive accuracy, achieving a Pearson correlation coefficient of 0.4741 and a MSE of 0.2029. R^2^ values of the test set were negative, indicating substantially reduced predictive performance on these synthetic variants. In a LOO-CV setting that incorporated both training and test data, the models also exhibited lower predictive performance compared to only trained with the training data, with RF achieving the best metrics (R^2^=0.3791, MSE=0.0890, Pearson correlation coefficient=0.6314) (Supplementary Table 14). Despite this drop in performance, nearly all data points, except for a single outlier, fell within the confidence interval of the GP models when assessing the correlation between true and predicted K_D_ values (Figure 5D). Our findings confirm that the decreased performance can largely be attributed to a single outlier within the test set, as the evaluation metrics, especially MSE, are particularly susceptible to outliers. When this data point is excluded, performance metrics improved substantially, with the GP_RBF model demonstrating the most accurate predictions, achieving an R^2^ value of 0.8669, Pearson correlation of 0.9378 and an MSE of 0.0236 (Supplementary Table 15).

**Figure 5.**
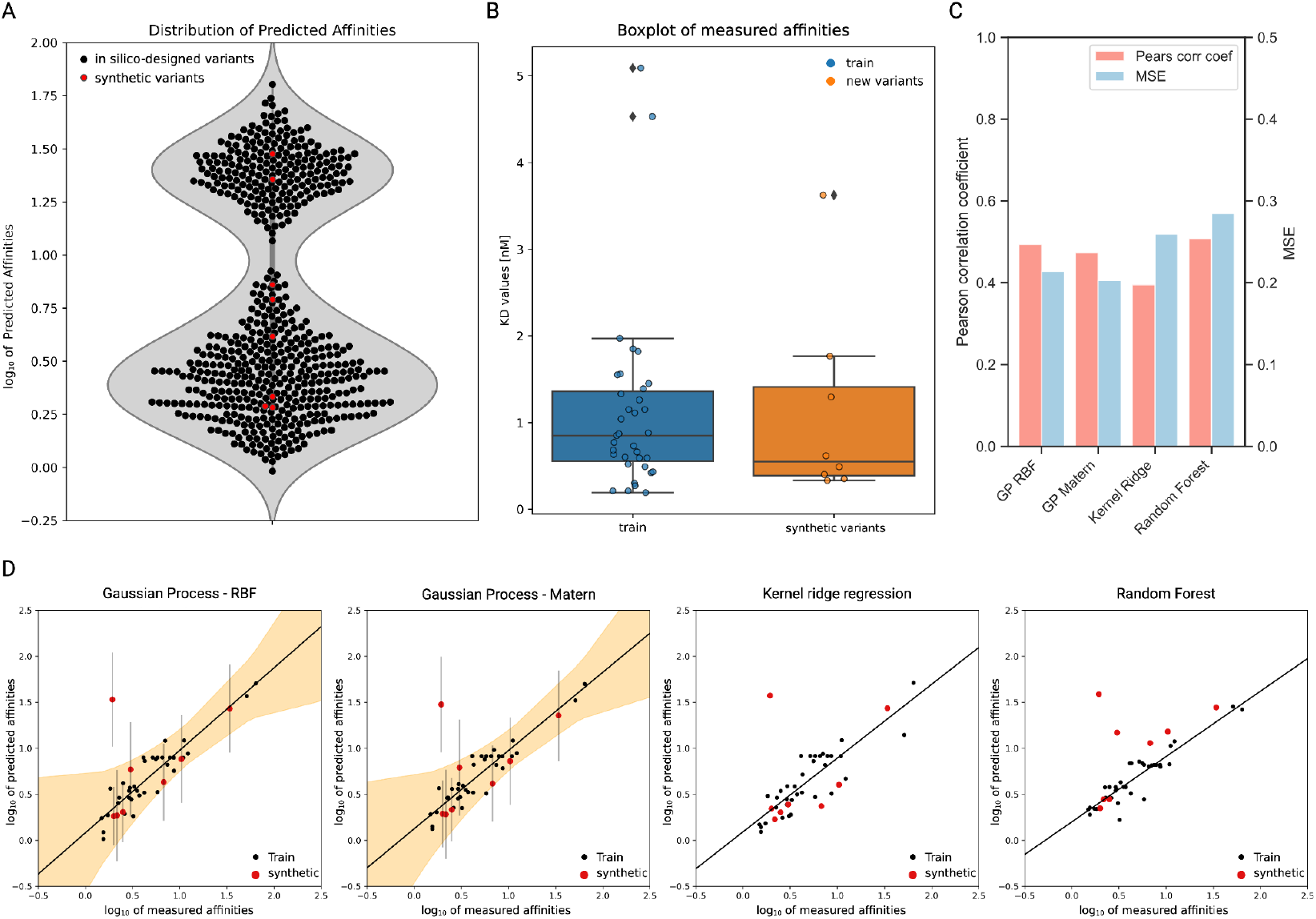
Experimental validation of synthetic (*in silico*-designed) VH variant antibodies. (**A**) Violin plot and swarm plot of the log_10_ predicted affinities of the synthetic antibody VH variants. The selected synthetic variants are depicted in red. (**B**) Boxplots displaying the experimentally determined K_D_ values of the training set and the synthetic variants. (**C**) Bar plots showing Pearson correlation coefficients and MSE of the trained ML models in predicting the synthetic variants. (**D**) Scatter plots illustrating the correlation between true and predicted values of the log10 of the training dataset (derived from LOO-CV) and the synthetic variants as a test set when the model was trained on the training dataset.

## Discussion

This work presents a case study of antibody engineering integrating antibody repertoire datasets, ML and experimental screening. An unsupervised, computational workflow was developed to identify antigen-specific variants from repertoire data that utilized a single known antigen-specific antibody (VH) sequence. 35 selected VH variants were expressed and their affinities were measured by BLI. ML models trained on the amino acid sequences of these variants achieved strong predictive performance for the measured affinities. To further evaluate the utility of these models for affinity engineering, synthetic antibody variants were designed *in silico*, guided by the ML predictions. Experimental characterization validated the predicted affinities for seven out of eight synthetic variants, highlighting the accuracy and reliability of this approach. Our work illustrates the potential of combining computational and experimental strategies to refine and engineer antibody affinity while minimizing the need for extensive experimental screening.

In order to generate training data for the ML models, we proposed a workflow for the selection of antigen-specific variants. Even though this workflow was able to identify multiple high affinity binders, we only considered antibody VH sequences of identical length (primarily determined by CDRH3) to the known initial binding variant. While this constraint primarily avoided the need to pad the sequences for the ML models, considering sequences of varying length may also reduce the probability of binding in the selected sequences. By applying this filtering the available antibody repertoire dataset was considerably reduced and otherwise potentially valuable variants were excluded solely based on sequence length. Developing a workaround that enables the inclusion of variable-length sequences, would allow for more comprehensive utilization of the antibody repertoire data. Such improvements could involve advanced encoding techniques, like variable-length sequence embeddings or PLM-derived embeddings (Marquet et al. 2022; Hie et al. 2023), to fully utilize the diversity of the available datasets. Despite this limitation, our proposed selection workflow highlights the potential of unsupervised methods for the discovery of additional binders, leveraging simple similarity-based approaches and antibody repertoire datasets.

The ML models demonstrated strong performance when evaluated with the training dataset, but the experimental validation results revealed a decrease in performance for out-of-distribution synthetic variants. Notably, this decrease in predictive capability was attributed to a single data point (tHC7, K_D_ = 0.33 nM, predicted K_D_ = 3.38 nM), which could be an outlier and may represent a synthetic variant with mutations that have more complex interactions contributing to its binding affinity, which are not adequately captured by the limited training data. Increasing training data, including expansion of the antibody sequence space may allow for a more detailed analysis of potential (non-)synergistic effects. Furthermore, employing more complex models, such as Bayesian Neural Networks (Lee 2000), could leverage a larger dataset and model sequence-affinity interactions more accurately, while still offering probabilistic outputs. Looking forward, GPs not only provide a reliable predictive framework, but also enable the application of a practical optimization approach such as Bayesian optimization (BO) (Garnett 2023). BO leverages the probabilistic output of GPs to suggest data points with optimal characteristics, predicted based on the available training data. This approach has demonstrated its potential in protein engineering studies (Romero et al. 2013; Hie et al. 2020; Li et al. 2023). By deploying BO, future studies could prioritize the selection of variants - synthetic as well as natural variants from antibody repertoires - with optimized antigen affinity, enabling iterative improvements. Additionally, more detailed analysis of the antibody repertoire, especially examining the relationship between predicted affinity and SHM or clonal expansion levels, could yield valuable insights into the generation of high-affinity binding antibodies in the immune system.

In conclusion, we show that antibody repertoire datasets represent a valuable source for the discovery of antigen-specific antibodies. Moreover, our findings highlight the potential to enhance the efficiency of antibody engineering through the integration of computational and experimental approaches.

## Acknowledgements

We would like to gratefully acknowledge the ETH Zurich D-BSSE Single-Cell Facility, including Dr. Mariangela Di Tacchio, Dr. Aleksandra Gumienny. Computing resources and support from ETH Zurich (Euler cluster) are gratefully acknowledged. This work has been supported by the Swiss National Science Foundation to S.T.R.

Figures in this manuscript were created with BioRender.com.

## Author contributions

Conceptualization, L.E., D.N., S.F, and S.T.R.; experiments, L.E., D.N.; computational analysis and machine learning implementation, L.E., S.F.; supervision, D.N., S.F., D.M. and S.T.R.; funding acquisition, S.T.R.; writing, L.E., D.M. and S.T.R.

## Methods

### Sequence selection

The antibody repertoire dataset from mice immunized with HEL that was used in this study, was generated as described in (Friedensohn et al. n.d.). Unique VH sequences were considered for the sequence selection workflow. The initial 3A-WT VH variant with confirmed antigen-specificity was discovered internally. The computational workflow to select antigen-specific sequences as training dataset was implemented in Python using open-source packages in the SciPy and scikit-learn ecosystem (Virtanen et al. 2020; Pedregosa et al. 2011).

#### Similarity filtering

In order to filter for similarity to the binder sequence, Levenshtein (edit) distance (LD) normalized by sequence length was calculated for each CDRH3 sequence of the natural repertoire and subtracted from 1. Sequences were taken to further selection if their similarity was higher than 0.8.

#### Affinity propagation (AP) clustering

AP clustering developed by (Frey & Dueck 2007) creates clusters by assigning values to data points (sequences) based on similarity (1 - LD). These values represent the suitability of one sequence being a cluster center while taking into account other values of sequence pairs. Input to algorithm was provided as a similarity matrix of all the CDRH3 sequences. As all sequences were equally suitable as exemplars, the preferences of the sequences were set to the median of the input similarities. Sequences that are clustered together with the binder sequence were taken to further selection.

#### k-medoids clustering

To cover most of the sequence space and to obtain the most representative sequences on VH-level, k-medoids clustering was performed on the remaining VDJ sequences in the data set. k-medoids clustering minimizes distances between data points and cluster centroids, whereas cluster centroids are actual data points (Kaufmann & Rousseeuw 01 1987). The number of cluster centroids to be found were defined to be 50.

### Preprocessing

The VH sequences were pre-processed to be expressed in the Pnp platform. Since the alignment of the VH sequences lacked the first 7 aa residues in the FR1 region, IMGT-Vquest was used to align the sequences to the IMGT reference database. The missing residues were replaced by the residues of the aligned germline. Throughout this preprocessing a sequencing error in one of the selected VH variants was detected and therefore, this variant was excluded. This reduced the full dataset size to 49 variants.

### Circular HDR donor (Plasmid vector) amplification

The full VH and VL sequences were ordered as custom plasmid vectors (Twist Bioscience). To generate sufficient amounts of plasmid for the hybridoma transfections, bacterial cultures and a Midi-prep was performed according to the manufacturer’s instructions (Zymo, CAT#: D4201).

### Hybridoma cell culture conditions

The Hybridoma cell lines were cultivated in high-glucose Dulbecco’s Modified Eagle Medium (Thermo, 11960-044) supplemented with 10% fetal bovine serum (Thermo, 16000-044), 100 U/ml Penicillin/Streptomycin (Thermo, 15140-122), 2 mM Glutamine (Sigma-Aldrich, G7513), 10 mM HEPES buffer (Thermo, 15630-056) and 50 μM 2-mercaptoethanol (Sigma-Aldrich, M3148). The cells were maintained in 5 ml of culture in T-25 flasks (TPP, 90026) in incubators at a temperature of 37°C and 5% CO2 and typically passaged every 48/72 h. A list of all PnP cell lines is provided in Supplementary Table 8.

### Hybridoma transfection

Prior to the transfection, the cells were prepared as follows: 10^6^ cells were isolated and centrifuged at 125 x G for 10 minutes, washed with Opti-MEM^®^ I Reduced Serum Medium (Thermo, 31985-062), and centrifuged again with the same parameters. The cells were resuspended in 100 µl of total volume of nucleofection mix, containing 15 µg of the circular HDR donor constructs (Twist Bioscience), 0.5 nmol Alt-R gRNA (IDT) diluted in SF buffer (per kit manufacturer guidelines). Transfections of the cells were performed in 100 μl single Nucleocuvettes™ with the 4D-Nucleofector™ System (Lonza) using the SF Cell Line 4D-Nucleofector^®^ X Kit L (Lonza, V4XC-2024) with the program CQ-104. All experiments performed utilize constitutive expression of Cas9 from Streptococcus pyogenes (SpCas9). Following the transfection, the cells were typically cultured in 1 ml of growth media in 24-well plates (Thermo, 142475).

### Flow cytometry sorting

Flow cytometry-based single cell isolation was performed 48 h after transfection using the BD FACS Aria III (BD Biosciences). Before labeling with fluorescently conjugated antibody, the cells were washed with PBS. Then the Hybridoma cells were incubated with the labeling antibody for 30 min on ice, protected from light, washed again with PBS and sorted. The labeling reagents and working concentrations are described in Supplementary Table 7. For cell numbers different from 5×10^5^, the antibody/antigen amount and incubation volume were adjusted proportionally. After single-cell sorting, the cells were recovered in 100 µl conditioned medium in flat bottom 96-well plates (Eppendorf, 0030730119). The clones were eventually expanded in 48-(Greiner Bio-One, 677 180; Thermo, 150687), 24- and 12-well plates (Thermo, 150628) and progressively moved into six-well plates (Corning, 3516; Thermo, 40675) and T-25 flasks eventually. All cell lines generated and utilized in this study are listed in Supplementary Table 8.

### Screening for antigen binding by ELISA

To screen for antigen-specificity of the variants, ELISA plates (Corning, 3590) were coated overnight with purified hen egg lysozyme (HEL, Sigma-Aldrich, 62971-10G-F) concentrated at 4 μg/ml in PBS overnight. Plates were then blocked with PBS supplemented with 2% m/V milk (AppliChem, A0830) and 0.05% V/V Tween-20 (Sigma Aldrich, P9416-50ML). After blocking, the plates underwent three washing steps with PBS supplemented with Tween-20 0.05% V/V (PBST). 50 μl supernatants from cell culture and positive controls were incubated as duplicates at RT for 1 h, followed by three washing steps with PBST. A secondary HRP-conjugated antibody specific for V_k_ light chain (rat monoclonal anti-mouse, Abcam, AB99617) concentrated at 0.7 μg/ml was used. ELISA detection was performed using a 1-Step Ultra TMB-ELISA Substrate Solution (Thermo, 34028) as the HRP substrate, reaction was terminated with H_2_SO_4_ (1 M). Absorbance at 450 nm was read with Infinite 200 PRO NanoQuant (Tecan).

### Affinity measurement by BLI

Following expansion of the monoclonal populations, supernatants of all cell culture variants were collected and filtered through a 0.20 μm filter (Sartorius, 16534-K). Prior to affinity measurements, the supernatants were concentrated by ultrafiltration using centrifugal filters (Merck, UFC910024) in order to increase the quality of the data. The measurements were then performed on an Octet RED96e (FortéBio) using anti-mouse capture sensors (FortéBio, 18-5088) according to the following procedure. Sensors were hydrated in conditioned media diluted 1 in 2 with kinetics buffer (FortéBio, 18-1105) for at least 10 minutes before conditioning through 4 cycles of regeneration consisting of 10 seconds incubation in 10 mM glycine, pH 1.52 and 10 seconds in kinetics buffer. Conditioned sensors were then loaded with 0 μg/mL (reference sensor) or hybridoma supernatant followed by blocking with human IgG (Rockland, 009-0102) at 50 μg/mL in kinetics buffer. After the blocking step, loaded sensors were equilibrated in kinetics buffer and incubated with either 0 nM (reference sample), 2 nM, 5 nM or 25 nM HEL protein. For the antigen dissociation step, sensors were incubated in kinetics buffer. Kinetics analysis was performed in the analysis software Data Analysis HT v11.0.0.50.

### ML regression model implementation and testing

All ML models were implemented in Python using open-source packages in the SciPy ecosystem (Virtanen et al. 2020; Pedregosa et al. 2011). Amino acid sequences were one-hot encoded to generate input features. Hereby, the amino acid sequence *s* of length *L* is defined by a binary vector *x*_*se*_ of length *L* times 20 (for 20 possible amino acids) indicating the presence (1) or absence (0) of an amino acid at the respective position. Measured K_D_ values were normalized to have mean zero and standard deviation one.

In regression, the problem is to infer the value of an unknown function *f*(*x*) at a novel point *x* given observations *y* at inputs *X*.

The hyperparameters the models were optimized with are the following:

- GP_RBF: ‘kernel’: [RBF(l) for l in np.logspace(-1, 1, 3)], ‘alpha’: [1e-10, 1e-3, 0.1]
- GP_Matern: ‘kernel’: [Matern(l) for l in np.logspace(-1, 1, 3)], ‘alpha’: [1e-10, 1e-3, 0.1]
- KRR: ‘alpha’: [1e-3, 0.1, 1.0, 10.0], ‘kernel’: [‘linear’, ‘rbf’, ‘polynomial’], ‘degree’: [2, 3], ‘gamma’: [0.1, 1.0, 10.0]
- RF: ‘n_estimators’: [10, 100, 200], ‘max_depth’: [2, 5, 10]}
- LR: ‘fit_intercept’: [True, False]

#### LOO-CV

During each CV cycle one antibody sequence (test sample) was left out from the dataset (training data). The training data was used to optimize hyperparameters and to train the model, whereas the normalized K_D_ value of the test sample was unknown to the GP model and predicted. Cycles were repeated until every sample was left out and predicted once.

Knowing the true values and the predictions of the test samples of all CV cycles, correlation plots including variances of the predictions were drawn. Lastly, to evaluate model performance, regression metrics (R^2^, MSE) were calculated.

#### Nested CV

A robust method used to assess the performance of a machine learning model and ensure that the model selection and hyperparameter tuning are properly evaluated. The procedure involves two loops: an outer loop and an inner loop. In the outer loop, the data is split into training and test sets multiple times (using k-fold cross-validation). For each training set in the outer loop, the inner loop further splits this training data into additional training and validation sets (again using k-fold cross-validation) to perform hyperparameter tuning and model selection. The best model selected from the inner loop is then trained on the outer loop’s training set and evaluated on the outer loop’s test set. This process is repeated for each split in the outer loop, and the performance metrics from each iteration are averaged to provide a more reliable estimate of the model’s performance and its ability to generalize to new data. The value k for inner and outer loops in our model evaluation was 5. To evaluate model performance, regression metrics (R^2^, Pearson correlation coefficient, MSE) were calculated.

### Design and affinity prediction of in silico-designed antibody variants

To evaluate model performance beyond the variants in the antibody repertoire dataset, eight new sequence variants were designed through an *in silico* mutagenesis approach. The mutational space was defined by seed sequences that included the original antigen-specific variant, 3A-WT, and the two highest- and lowest-affinity variants from the training data. The *in silico*-designed variants were limited to single or double point mutations of the seed sequences. Out of all possible *in silico* variants eight sequences that span the affinity range as predicted by the trained GP_RBF model, were selected. These new *in silico*-designed sequences were then expressed using the PnP platform and characterized by BLI as described above.

