## Supplementary Information for "Antibody affinity engineering using antibody repertoire data and machine learning"

### Supplementary for GP publication

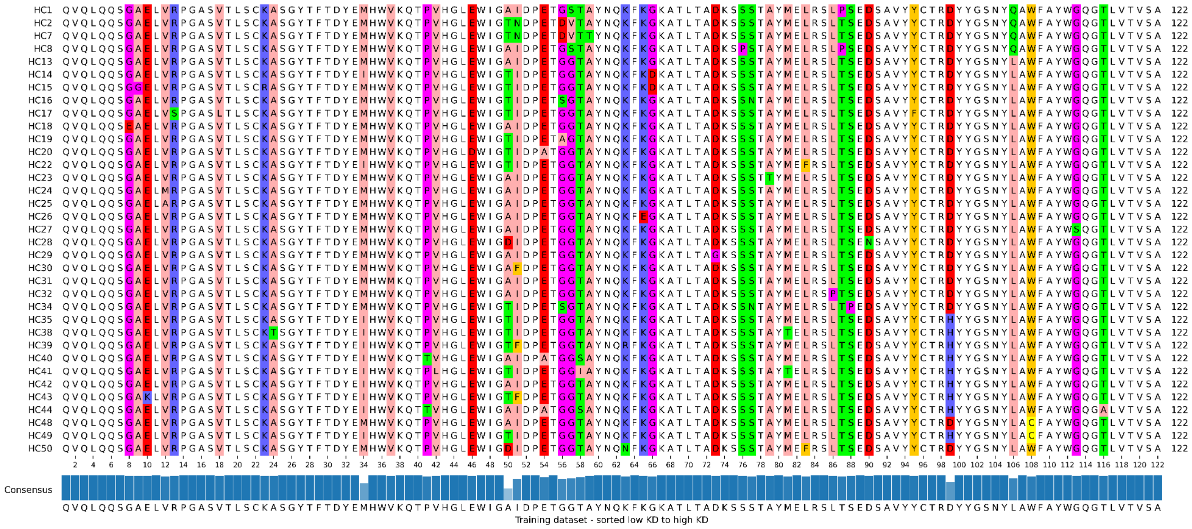

**Supplementary Figure 1. Amino acid sequence alignment of selected VDJ sequences.** Alignment of the selected VDJ sequences that were characterized experimentally. Amino acid positions with mutations are highlighted. The barplot in the bottom depicts the consensus sequence and identity bar.

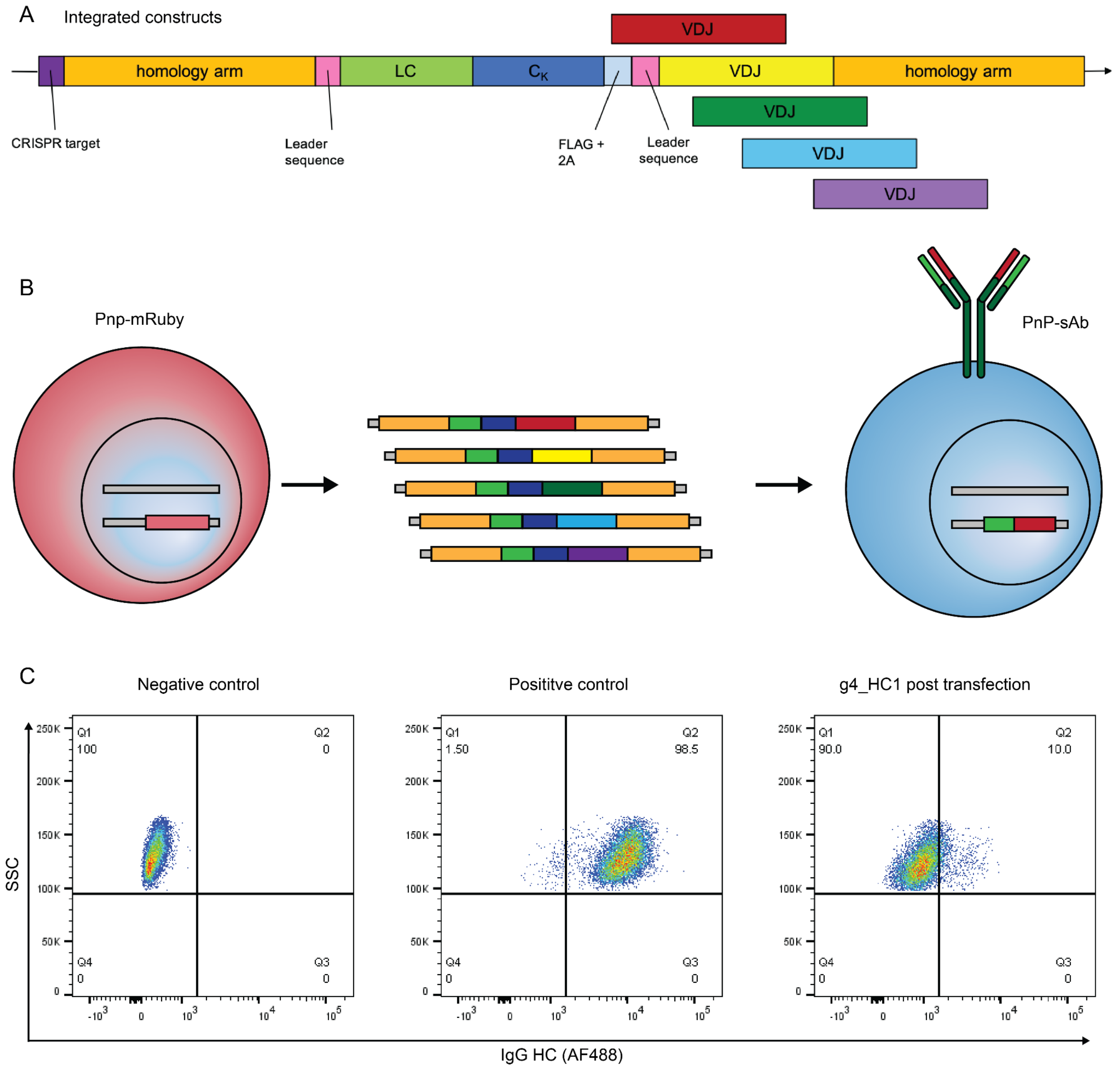


**Supplementary Figure 2. Experimental overview of cell line generation.** (**A**) Scheme of the gene constructs that were integrated into the genome of the Pnp-mRuby cell line to generate a cell line that expresses IgG. For details of the gene construct design, we refer to (Mason et al. 2018). (**B**) Schematics of the genomic integration into Pnp-mRuby cell line generated Pnp cell lines that express full IgG molecules on their surface. (**C**) Exemplary plots of the FC signal (IgG HC, AF488 vs. side scatter (SSC)). In the left and middle panel a $IgG^-$ cell line as negative control and an $IgG^+$ as positive control are depicted. In the right panel is the fluorescent signal of a transfected Pnp cell line, g4_HC1, 2 days post transfection with the gene constructs. Q2 gate was sorted as single cells for expansion.

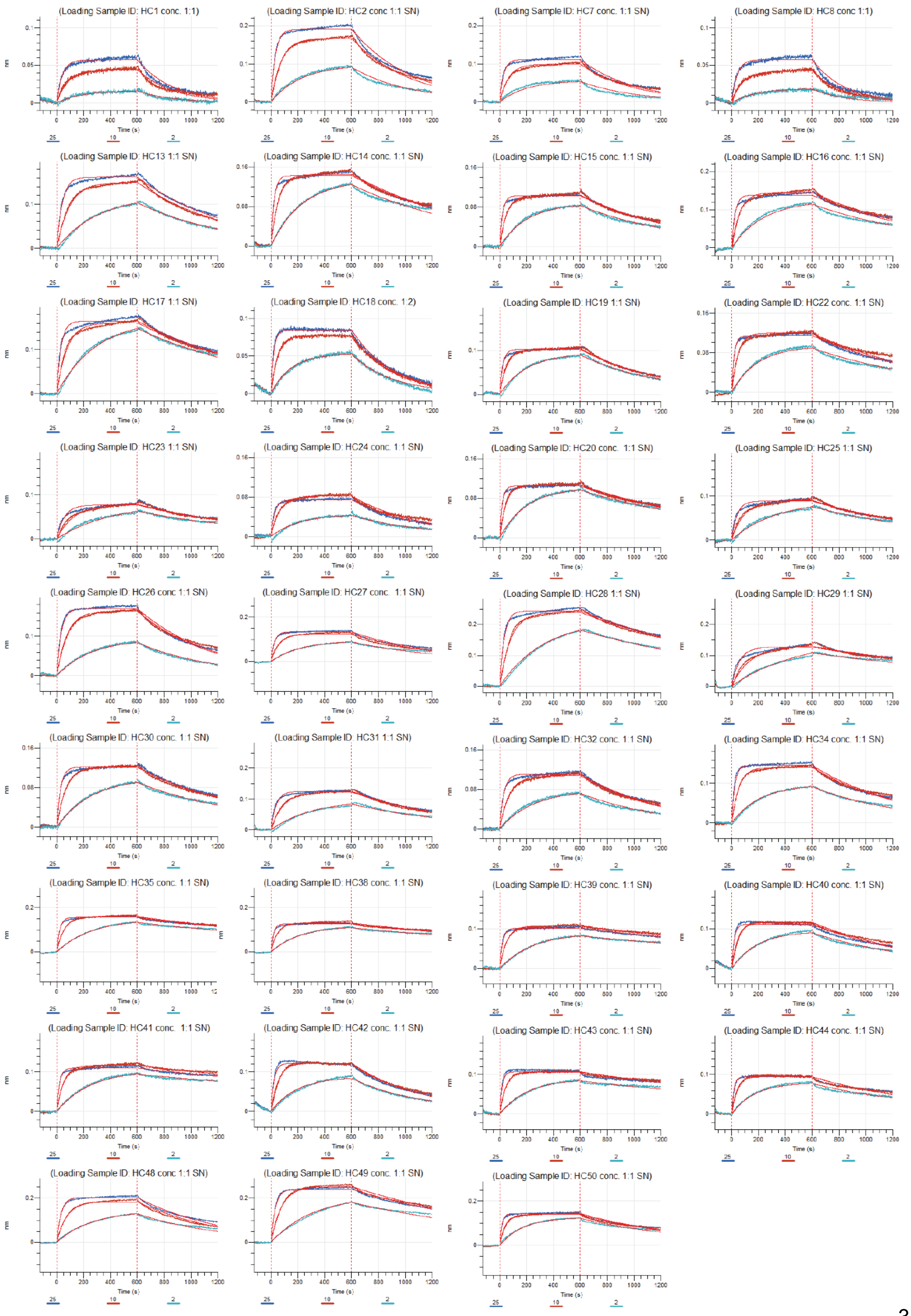
(Loading Sample ID: HC1 conc. 1:1)
(Loading Sample ID: HC2 conc 1:1 SN)
(Loading Sample ID: HC7 conc. 1:1 SN)
(Loading Sample ID: HC8 conc 1:1)
(Loading Sample ID: HC13 1:1 SN)
(Loading Sample ID: HC14 conc. 1:1 SN)
(Loading Sample ID: HC15 conc. 1:1 SN)
(Loading Sample ID: HC16 conc. 1:1 SN)
(Loading Sample ID: HC17 1:1 SN)
(Loading Sample ID: HC18 conc. 1:2)
(Loading Sample ID: HC19 1:1 SN)
(Loading Sample ID: HC22 conc. 1:1 SN)
(Loading Sample ID: HC23 1:1 SN)
(Loading Sample ID: HC24 conc. 1:1 SN)
(Loading Sample ID: HC20 conc. 1:1 SN)
(Loading Sample ID: HC25 1:1 SN)
(Loading Sample ID: HC26 conc 1:1 SN)
(Loading Sample ID: HC27 conc. 1:1 SN)
(Loading Sample ID: HC28 1:1 SN)
(Loading Sample ID: HC29 1:1 SN)
(Loading Sample ID: HC30 conc. 1:1 SN)
(Loading Sample ID: HC31 1:1 SN)
(Loading Sample ID: HC32 conc. 1:1 SN)
(Loading Sample ID: HC34 conc. 1:1 SN)
(Loading Sample ID: HC35 conc. 1:1 SN)
(Loading Sample ID: HC38 conc. 1:1 SN)
(Loading Sample ID: HC39 conc. 1:1 SN)
(Loading Sample ID: HC40 conc. 1:1 SN)
(Loading Sample ID: HC41 conc. 1:1 SN)
(Loading Sample ID: HC42 conc. 1:1 SN)
(Loading Sample ID: HC43 conc. 1:1 SN)
(Loading Sample ID: HC44 conc. 1:1 SN)
(Loading Sample ID: HC48 conc 1:1 SN)
(Loading Sample ID: HC49 conc. 1:1 SN)
(Loading Sample ID: HC50 conc. 1:1 SN)
Time (s)
nm
25
10
2

**Supplementary Figure 3. BLI binding curves.** Binding curves of the variants selected from immune repertoire that could be expressed sufficiently were measured by BLI of the variants selected from immune repertoire datasets. The graphs depict binding, measured as nm, over time.

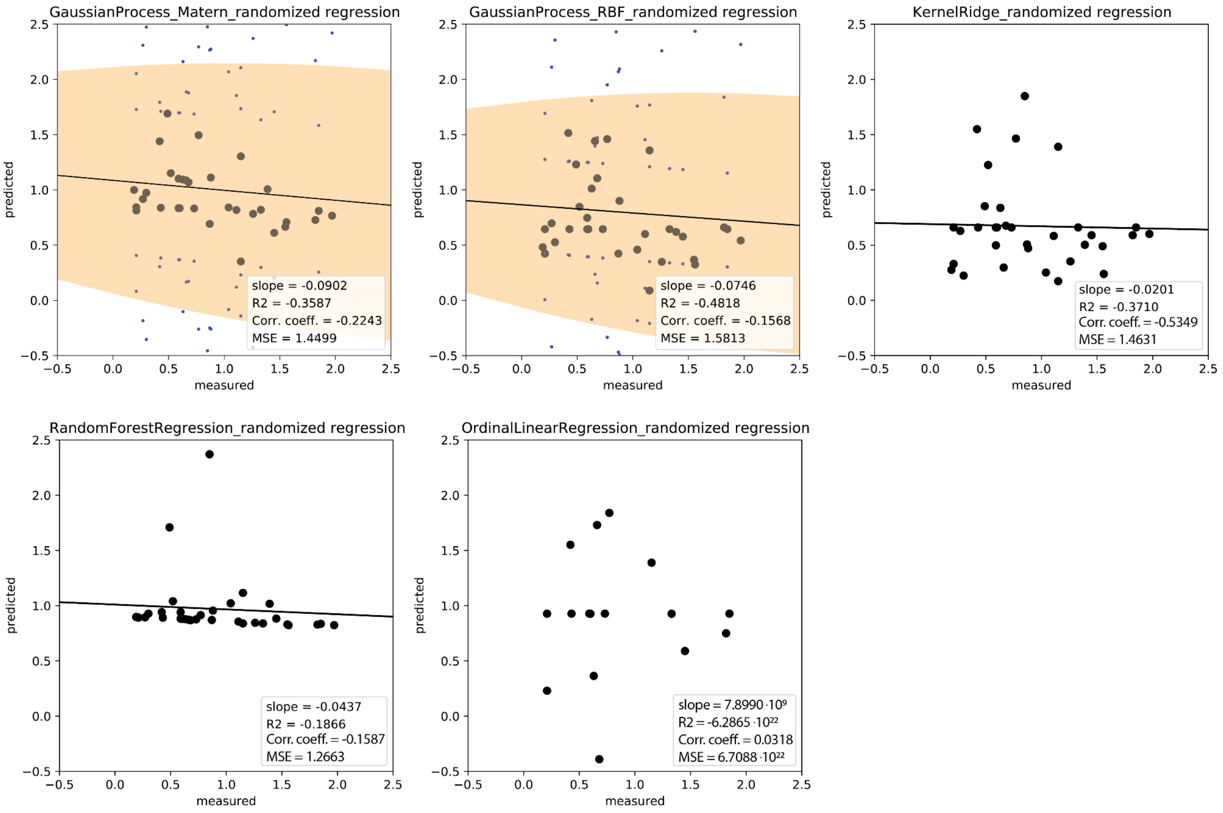


**Supplementary Figure 4. Correlation plots of the ML models trained on the training data with randomized labels.** Each dot in the scatter plots represents the predicted vs measured KD value. Predictions were obtained from LOO-CV. Slope, $R^2$, Pearson correlation coefficient and MSE metrics of each respective model in the legends.

**Supp. Table 1. Immunization scheme**

| Group | n= 3 | Primary | | 1st boost | | 2nd boost | | 3rd boost |
|---|---|---|---|---|---|---|---|---|
| | | Antigen | Adjuvant | Antigen | Adjuvant | Antigen | Adjuvant | Antigen |
| **HEL, 3 boosts** | 3 | 200 µg HEL | 20 µg MPLA | 50 µg HEL | 20 µg MPLA | 50 µg HEL | 20 µg MPLA | 50 µg HEL |
| **HEL, 2 boosts** | 3 | 200 µg HEL | 20 µg MPLA | 50 µg HEL | 20 µg MPLA | 50 µg HEL | | |
| **HEL, 1 boosts** | 3 | 200 µg HEL | 20 µg MPLA | 50 µg HEL | | | | |
| **HEL, no boosts** | 3 | 200 µg HEL | 20 µg MPLA | | | | | |

**Supp. Table 2. Details of the known binder sequence 3A**

| | **known Binder (3A)** |
|---|---|
| **VDJ seq (len)** | GAELVRPGASVTLSCKASGYTFTDYEMHWVKQTPVHGLEWIGDIDPETGGTAYNQNFKGKATLTADKSSSTAYMEFRSLTSEDSAVYYCTRDYYGSNYLAWFAYWGQGTLVTVSA (115) |
| **CDRH3 seq (len)** | CTRDYYGSNYLAWFAYW (17) |
| **VGene** | IGHV1-15*01 |
| **JGene** | IGHJ3*01 |
| **# SHM** | 4 |
| **Isotype** | mIgG1 |

**Supp. Table 3. Number of antibody sequences in the dataset**

| | Number of sequences in antibody repertoire |
|---|---|
| **unique VDJs** | 83478 |
| **unique CDRH3s** | 27076 |
| **VDJs (115 amino acids long)** | 4610 |
| **CDRH3 (17 amino acids long)** | 1761 |

**Supp. Table 4. Repertoire stats of the 193 VDJ sequences from CDRH3 selection.**

| | Repertoire stats of the selected VDJ sequences |
|---|---|
| **VGenes (and %)** | IGHV1-15*01 (92.7%)<br>IGHV1-69*02 (1.6%)<br>IGHV1-20*02 (1.0%)<br>IGHV1-7*01 (1.0%)<br>IGHV1-67*01 (0.5%)<br>IGHV1-80*01 (0.5%)<br>IGHV1-4*01 (0.5%)<br>IGHV1-9*01 (0.5%)<br>IGHV1-77*01 (0.5%)<br>IGHV1-39*01 (0.5%)<br>IGHV1-52*01 (0.5%) |
| **JGenes (and %)** | IGHJ3*01 (100.0%) |
| **Isotypes (and %)** | mIgG1 (97.9%)<br>NoIsotype (1.6%)<br>mIgG2b (0.5%) |
| **mean # of SHMs (min, max)** | 4.2 (1, 19) |

**Supp. Table 5. Sequences of all selected/ordered antibody variants**

| SampleID | Sequence |
|---|---|
| HC1 | QVQLQQSGAELVRPGASVTLSCKASGYTFTDYEMHWVKQTPVHGLEWIGAIDPETGSTAYNQKFKGKAT<br>LTADKSSSTAYMELRSLPSEDSAVYYCTRDYYGSNYQAWFAYWGQGTLVTVSA |
| HC2 | QVQLQQSGAELVRPGASVTLSCKASGYTFTDYEMHWVKQTPVHGLEWIGTNDPETDVTAYNQKFKGKA<br>TLTADKSSSTAYMELRSLTSEDSAVYYCTRDYYGSNYQAWFAYWGQGTLVTVSA |
| HC3 | QVQLQQSGAELVRPGASVTLSCKASGYTFTDYEMHWVKQTPVHGLEWIGAIDPETGGTAYNQKFKGKAT<br>LTADKSSSTAYMELRSLTSEDSAVYYCTRDYYGSNYQAWFAYWGQGTLVTVSA |
| HC4 | QVQLQQSGAELVRPGASVTLSCKASGYTFTDYEMHWVKQTPVHGLEWIGAIDPETGSTAYNQKFKGKAT<br>LTADKSSSTAYMELRSLPSEVSAVYYCTRDYYGSNYQAWFAYWGQGTLVTVSA |
| HC5 | QVQLQQSGAELVRPGASVTLSCKASGYTFTDYEMHWVKQTPVHGLEWIGAIDPKTGSTAYNQKFKGKAT<br>LTADKSSSTAYMELRSLPSEDSAVYYCTRDYYGSNYQAWFAYWGQGTLVTVSA |
| HC7 | QVQLQQSGAELVRPGASVTLSCKASGYTFTDYEMHWVKQTPVHGLEWIGTNDPETDVTTYNQKFKGKA<br>TLTADKSSSTAYMELRSLTSEDSAVYYCTRDYYGSNYQAWFAYWGQGTLVTVSA |
| HC8 | QVQLQQSGAELVRPGASVTLSCKASGYTFTDYEMHWVKQTPVHGLEWIGAIDPETGSTAYNQKFKGKAT<br>LTADKSPSTAYMELRSLPSEDSAVYYCTRDYYGSNYQAWFAYWGQGTLVTVSA |
| HC9 | EVQLQQSGPELVKPGASVKISCKASGYSFTGYFMNWVKQSHGKSLEWIGCINPYNGDTFYNQKFKGKAT<br>LTADKSSSTAYMELRSLPSEDSAVYYCTRDYYGSNYQAWFAYWGQGTLVTVSA |
| HC10 | QVQLQQSGAELVRPGASVTLSCKASGYTFTDYEMHWVKQTPVHGLEWIGAIDPETGSTAYNQKFKGKAT<br>LTADKSSSTAYMELRSLPSEDSAVDYCTRDYYGSNYQAWFAYWGQGTLVTVSA |
| HC11 | QVQLQQSGAELAKPGASVKMSCKASGYTFTSYWMHWVKQRPGQGLEWIGYINPSTSYTEYNQKFKDK<br>ATLTADKSSSTAYMELRSLPSEDSAVYYCTRDYYGSNYQAWFAYWGQGTLVTVSA |
| HC12 | QVQLQQSGAELVRPGASVTLSCKASGYTFTDYEIHWVKQTPVHGLEWIGAIDPETDGTAYNQKFKGKATL<br>TADKSSNTAYMELRSLTSEDSAVYYCTRDYYGSSYQAWFTYWGQGTLVTVSA |
| HC13 | QVQLQQSGAELVRPGASVTLSCKASGYTFTDYEMHWVKQTPVHGLEWIGAIDPETGGTAYNQKFKGKAT<br>LTADKSSSTAYMELRSLTSEDSAVYYCTRDYYGSNYLAWFAYWGQGTLVTVSA |
| HC14 | QVQLQQSGAELVRPGASVTLSCKASGYTFTDYEIHWVKQTPVHGLEWIGTIDPETGGTAYNQKFKDKATL<br>TADKSSSTAYMELRSLTSEDSAVYYCTRDYYGSNYLAWFAYWGQGTLVTVSA |
| HC15 | QVQLQQSGGELVRPGASVTLSCRASGYTFTDYEMHWVKQTPVHGLEWIGTIDPETGGTAYNQKFKDKAT<br>LTADKSSSTAYMELRSLTSEDSAVYYCTRDYYGSNYLAWFAYWGQGTLVTVSA |
| HC16 | QVQLQQSGAELVRPGASVTLSCKASGYTFTDYEMHWVKQTPVHGLEWIGTIDPETSGTAYNQKFKGKAT<br>LTADKSSNTAYMELRSLTSEDSAVYYCTRDYYGSNYLAWFAYWGQGTLVTVSA |
| HC17 | QVQLQQSGAELVSPGASLTLSCKASGYTFTDYEMHWVKQTPVHGLEWIGTIDPETGGTAYNQKFKGKAT<br>LTADKSSSTAYMELRSLTSEDSAVYFCTRDYYGSNYLAWFAYWGQGTLVTVSA |
| HC18 | QVQLQQSEAELVRPGASVTLSCKASGYTFTDYEMHWVKQTPVHGLEWIGAIDPETGGTAYNQKFKGKAT<br>LTADKSSSTAYMELRSLTSEDSAVYYCTRDYYGSNYLAWFAYWGQGTLVTVSA |
| HC19 | QVQLQQSGAELVRPGASVTLSCKASGYTFTDYEMHWVKQTPVHGLEWIGTIDPETAGTAYNQKFKGKAT<br>LTADKSSSTAYMELRSLTSEDSAVYYCTRDYYGSNYLAWFAYWGQGTLVTVSA |
| HC20 | QVQLQQSGAELVRPGASVTLSCKASGYTFTDYEMHWVKQTPVHGLDWIGTIDPATGGTAYNQKFKGKA<br>TLTADKSSSTAYMELRSLTSEDSAVYYCTRDYYGSNYLAWFAYWGQGTLVTVSA |
| HC21 | QVQLQQSGAELVRPGASVTLSCKASGYTFTDYEVHWVKQTPVHGLEWIGTIDPETGNTAYNQKFKGKAT<br>LTADKSSSTAYMEFRSLTSEDSAVCYCTRDYYGSNYLAWFAYWGQGTLVTVSA |
| HC22 | QVQLQQSGAELVRPGASVTLSCKASGYTFTDYEIHWVKQTPVHGLEWIGTIDPETGGTAYNQKFKGKATL<br>TADKSSSTAYMEFRSLTSEDSAVYYCTRDYYGSNYLAWFAYWGQGTLVTVSA |
| HC23 | QVQLQQSGAELVRPGASVTLSCKASGYTFTDYEMHWVKQTPVHGLEWIGAIDPETGGTAYNQKFKGKAT<br>LTADKSSSTTYMELRSLTSEDSAVYYCTRDYYGSNYLAWFAYWGQGTLVTVSA |
| HC24 | QVQLQQSGAELMRPGASVTLSCKASGYTFTDYEMHWVKQTPVHGLEWIGAIDPETGGTAYNQKFKGKA<br>TLTADKSSSTAYMELRSLTSEDSAVYYCTRDYYGSNYLAWFAYWGQGTLVTVSA |
| HC25 | QVQLQQSGAELARPGASVTLSCKASGYTFTDYEMHWVKQTPVHGLEWIGAIDPETGGTAYNQKFKGKA<br>TLTADKSSSTAYMELRSLTSEDSAVYYCTRDYYGSNYLAWFAYWGQGTLVTVSA |
| HC26 | QVQLQQSGAELVRPGASVTLSCKASGYTFTDYEMHWVKQTPVHGLEWIGAIDPETGGTAYNQKFEGKAT<br>LTADKSSSTAYMELRSLTSEDSAVYYCTRDYYGSNYLAWFAYWGQGTLVTVSA |
| HC27 | QVQLQQSGAELVRPGASVTLSCKASGYTFTDYEMHWVKQTPVHGLEWIGAIDPETGGTAYNQKFKGKAT<br>LTADKSSSTAYMELRSLTSEDSAVYYCTRDYYGSNYLAWFAYWSQGTLVTVSA |
| HC28 | QVQLQQSGAELVRPGASVTLSCKASGYTFTDYEMHWVKQTPVHGLEWIGDIDPETGGTAYNQKFKGKA<br>TLTADKSSSTAYMELRSLTSENSAVYYCTRDYYGSNYLAWFAYWGQGTLVTVSA |
| HC29 | QVQLQQSGAELVRPGASVTLSCKASGYTFTDYEMHWVKQTPVHGLEWIGAIDPETGGTAYNQKFKGKAT<br>LTAGKSSSTAYMELRSLTSEDSAVYYCTRDYYGSNYLAWFAYWGQGTLVTVSA |

| | |
|---|---|
| HC30 | QVQLQQSGAELVRPGASVTLSCKASGYTFTDYEMHWVKQTPVHGLEWIGAFDPETGGTAYNQKFKGKA TLTADKSSSTAYMELRSLTSEDSAVYYCTRDYYGSNYLAWFAYWGQGTLVTVSA |
| HC31 | QVQLQQSGAELVRPGASVTLSCKASGYTFTDYEMHWMKQTPVHGLEWIGAIDPETGGTAYNQKFKGKA TLTADKSSSTAYMELRSLTSEDSAVYYCTRDYYGSNYLAWFAYWGQGTLVTVSA |
| HC32 | QVQLQQSGAELVRPGASVTLSCKASGYTFTDYEMHWVKQTPVHGLEWIGAIDPETGGTAYNQKFKGKAT LTADKSSSTAYMELRSPTSEDSAVYYCTRDYYGSNYLAWFAYWGQGTLVTVSA |
| HC33 | QVQLQQSGAELVRPGVSVKISCKGSGYTFTDYAIHWVKQSHAKSLEWIGVISTYYSDTTYNQKFKGKATM TADKSSSTAYMELRSLTSEDSAVYYCTRDYYGSNYLAWFAYWGQGTLVTVSA |
| HC34 | QVQLQQSGAELVRPGASVTLSCKASGYTFTDYEMHWVKQTPVHGLEWIGTIDPETSGTAYNQKFKGKAT LTADKSSNTAYMELRSLTPEDSAVYYCTRDYYGSNYLAWFAYWGQGTLVTVSA |
| HC35 | QVQLQQSGAELVRPGASVTLSCKASGYTFTDYEIHWVKQTPVHGLEWIGTIDPETGGTAYNQKFKGKATL TADKSSSTAYMELRSLTSEDSAVYYCTRHYYGSNYLAWFAYWGQGTLVTVSA |
| HC36 | QVQLQQSGAELVRPGASVTLSCKASGYTFTDYEIHWVKQTPVHGLEWIGTIDPETDSTAYNQKFKGKATL TADKSSSTAYMELRSLTSEDSAVYYCTRHYYGSNYLAWFAYWGQGTLVTVSA |
| HC37 | QVQLQQSGAELVRPGASVTLSCKASGYTFTDYEIHWVKQTPVHGLEWIGTLDPETDDTAYSQKFKGKATL TADKSSSTAYMELRSLTSEDSAVYYCTRHYYGSNYLAWFAYWGQGTLVTVSA |
| HC38 | QVQLQQSGAELVRPGASVTLSCKTSGYTFTDYEIHWVKQTPVHGLEWIGTIDPETGGTAYNQKFKGKATL TADKSSSTAYTELRSLTSEDSAVYYCTRHYYGSNYLAWFAYWGQGTLVTVSA |
| HC39 | QVQLQQSGAELVRPGASVTLSCKASGYTFTDYEIHWVKQTPVHGLEWIGTFDPETGGTAYNQRFKGKAT LTADKSSSTAYMELRSLTSEDSAVYYCTRHYYGSNYLAWFAYWGQGTLVTVSA |
| HC40 | QVQLQQSGAELVRPGASVTLSCKASGYTFTDYEIHWVKQTTVHGLEWIGAIDPATGGSAYNQKFKGKATL TADKSSSTAYMELRSLTSEDSAVYYCTRHYYGSNYLAWFAYWGQGTLVTVSA |
| HC41 | QVQLQQSGAELVRPGASVTLSCKASGYTFTDYEIHWVKQTPLHGLEWIGTIDPETGGIAYNQKFKGKATL TADKSSSTAYTELRSLTSEDSAVYYCTRHYYGSNYLAWFAYWGQGTLVTVSA |
| HC42 | QVQLQQSGAELVRPGASVTLSCKASGYTFTDYEIHWVKQTPVHGLEWIGAIDPETGGTAYNQKFKGKATL TADKSSSTAYMELRSLTSEDSAVYYCTRHYYGSNYLAWFAYWGQGTLVTVSA |
| HC43 | QVQLQQSGAKLVRPGASVTLSCKASGYTFTDYEIHWVKQTPVHGLEWIGTFDPETGGTAYNQKFKGKAT LTADKSSSTAYMELRSLTSEDSAVYYCTRHYYGSNYLAWFAYWGQGTLVTVSA |
| HC44 | QVQLQQSGAELVRPGASVTLSCKASGYTFTDYEIHWVKQTTVHGLEWIGAIDPATGGSAYNQKFKGKATL TADKSSSTAYMELRSLTSEDSAVYYCTRHYYGSNYLAWFAYWGQGALVTVSA |
| HC45 | QVQLQQPGAELVRPGASVKLSCKASGYTFTSYWINWVRQRPGQGLEWIGNIYPSDSYTNYNQKFKDKA TLTVDKSSSTAYMQLRSLTSEDSAVYYCTRHYYGSNYLAWFAYWGQGTLVTVSA |
| HC46 | QVQLQQSGAELVRPGASVTLSCKASGYTFTDYEMHWVKQTPVHGLEWIGDIDPETGGTAYNQKFKGKA TLTADKSSSTAYMELRSLTSEDSAVYYCTRDYYGSSYLAWFAYWGQGTLVTVSA |
| HC47 | QVQLQQSGAELVRPGASVTLSCKASGYTFTDYEMHWVKQTPVHGLEWIGTIDPETDGTAYNQKFKGKAT LTADKSSSTAYMELRSLTSEDSAVYYCTRGYYGSNYLAWFAYWGQGTLVTVSA |
| HC48 | QVQLQQSGAELVRPGASVTLSCKASGYTFTDYEMHWVKQTPVHGLEWIGAIDPETGGTAYNQKFKGKAT LTADKSSSTAYMELRSLTSEDSAVYYCTRDYYGSNYLACFAYWGQGTLVTVSA |
| HC49 | QVQLQQSGAELVRPGASVTLSCKASGYTFTDYEIHWVKQTPVHGLEWIGTIDPETGGTAYNQKFKGKATL TADKSSSTAYMELRSLTSEDSAVYYCTRHYYGSNYLACFAYWGQGTLVTVSA |
| HC50 | QVQLQQSGAELVRPGASVTLSCKASGYTFTDYEMHWVKQTPVHGLEWIGDIDPETGGTAYNQNFKGKA TLTADKSSSTAYMEFRSLTSEDSAVYYCTRDYYGSNYLAWFAYWGQGTLVTVSA |

**Supp. Table 6. Repertoire stats of the VDJ sequences from CDR3 selection**

| | Repertoire stats of the 50 selected VDJ sequences |
|---|---|
| **VGenes (and %)** | IGHV1-15*01 (92%)<br>IGHV1-67*01 (2%)<br>IGHV1-69*02 (2%)<br>IGHV1-20*02 (2%)<br>IGHV1-7*01 (2%) |
| **JGenes (and %)** | IGHJ3*01 (100.0%) |
| **Isotypes (and %)** | mIgG1 (98%)<br>mIgG2b (2%) |
| **mean # of SHMs (min, max)** | 4 (1, 10) |

**Supp. Table 7. FC labelling reagents and working concentrations**

| Antibody/ Antigen | Working conc. | Dilution from stock | Incubation volume | Conjugated Fluorophore | Product Id |
|---|---|---|---|---|---|
| **Anti-IgG2C**[12] | 12 µg/ml | 1:100 | 50 ul | AlexaFluor® 488 | 115-545-208 (Jackson ImmunoResearch) |
| **Anti-IgK**[2] | 2.5 µg/ml | 1:80 | 50 µl | Brilliant Violet 421™ | 409511 (BioLegend) |
| **HEL** [2] | 0.99 µg/ml | 1:62.5 | 50 µl | AlexaFluor® 647 | 62971-10G-F (Sigma- Aldrich) |

[1] used for single-cell sort
[2] used for for cytometric analysis

**Supp. Table 8. List of all PnP cell lines used**

| Cell Name | Cell Name (Abrev.) | Description |
|---|---|---|
| **PnP-mRuby-Cas9** | RC9 | Starting cell line for generation of g4_HC cell lines; hybridoma cell line used as antibody display and expression system; native light chain locus was deleted and native heavy chain locus was replaced by mRuby reporter gene; cell line was also engineered to constitutively express Cas9 from Rosa26 safe harbour locus |
| **PnP-HEL23** | Y | Hybridoma cell line immunogenomically engineered to produce antigen specific antibodies targeting hen egg lysozyme (HEL). The light and heavy chains are expressed from a single transcript containing a 2A, self-cleaving peptide. |
| **PnP-HC1-50** | g4_HC1-50 | RC9 hybridoma cell line immunogenomically engineered to produce antigen specific antibodies targeting hen egg lysozyme (HEL). The light and heavy chains are expressed from a single transcript containing a 2A, self-cleaving peptide. |

**Supp. Table 9. BLI affinity measurements of the training dataset**

| Sample | $K_D$ value [nM] | $k_a$ [$M^{-1}s^{-1}$] | $k_d$ [$s^{-1}$] | $R^2$ |
|---|---|---|---|---|
| HC41 | 0.189 | 1.90E+06 | 3.59E-04 | 0.9840 |
| HC39 | 0.211 | 1.91E+06 | 4.03E-04 | 0.9917 |
| HC43 | 0.215 | 2.26E+06 | 4.87E-04 | 0.9693 |
| HC38 | 0.269 | 2.08E+06 | 5.58E-04 | 0.9766 |
| HC35 | 0.299 | 1.75E+06 | 5.24E-04 | 0.9830 |
| HC20 | 0.418 | 2.12E+06 | 8.87E-04 | 0.9833 |
| HC44 | 0.427 | 2.49E+06 | 1.06E-03 | 0.9640 |
| HC50 | 0.491 | 2.42E+06 | 1.19E-03 | 0.9667 |
| HC22 | 0.519 | 2.08E+06 | 1.08E-03 | 0.9789 |
| HC19 | 0.588 | 1.68E+06 | 1.64E-03 | 0.9878 |
| HC40 | 0.590 | 2.09E+06 | 1.23E-03 | 0.9785 |
| HC28 | 0.602 | 1.17E+06 | 7.01E-04 | 0.9912 |
| HC14 | 0.631 | 1.68E+06 | 1.06E-03 | 0.9825 |
| HC49 | 0.660 | 1.21E+06 | 8.00E-04 | 0.9882 |
| HC16 | 0.680 | 1.56E+06 | 1.12E-03 | 0.9743 |
| HC15 | 0.726 | 1.82E+06 | 1.32E-03 | 0.9836 |
| HC17 | 0.765 | 1.23E+06 | 9.69E-04 | 0.9788 |
| HC29 | 0.854 | 6.66E+05 | 5.69E-04 | 0.9586 |
| HC34 | 0.872 | 1.70E+06 | 1.48E-03 | 0.9867 |
| HC30 | 0.885 | 1.36E+06 | 1.20E-03 | 0.9902 |
| HC27 | 1.037 | 1.52E+06 | 1.57E-03 | 0.9793 |

| | | | | |
|---|---|---|---|---|
| HC25 | 1.110 | 9.66E+05 | 1.07E-03 | 0.9690 |
| HC42 | 1.145 | 1.69E+06 | 1.94E-03 | 0.9889 |
| HC32 | 1.146 | 1.27E+06 | 1.46E-03 | 0.9860 |
| HC31 | 1.261 | 1.00E+06 | 1.26E-03 | 0.9863 |
| HC48 | 1.332 | 1.20E+06 | 1.60E-03 | 0.9883 |
| HC24 | 1.391 | 1.37E+06 | 1.91E-03 | 0.9849 |
| HC23 | 1.450 | 6.81E+05 | 9.87E-04 | 0.9579 |
| HC26 | 1.553 | 1.14E+06 | 1.77E-03 | 0.9907 |
| HC13 | 1.555 | 9.18E+05 | 1.43E-03 | 0.9917 |
| HC18 | 1.818 | 1.80E+06 | 3.28E-03 | 0.9944 |
| HC7 | 1.846 | 1.31E+06 | 2.43E-03 | 0.9757 |
| HC2 | 1.972 | 1.14E+06 | 2.25E-03 | 0.9891 |
| HC1 | 4.526 | 8.27E+05 | 3.74E-03 | 0.9809 |
| HC8 | 5.087 | 8.20E+05 | 4.17E-03 | 0.9781 |
| HC4 | negative ELISA | | | not expressed |
| HC10 | negative ELISA | | | not expressed |
| HC12 | negative ELISA | | | not expressed |
| HC37 | negative ELISA | | | not expressed |
| HC45 | negative ELISA | | | not expressed |
| HC3 | low quality BLI | | | not binding |
| HC33 | low quality BLI | | | not binding |
| HC36 | low quality BLI | | | not binding |
| HC47 | low quality BLI | | | not binding |
| HC5 | not binding (BLI) | | | not binding |
| HC9 | not binding (BLI) | | | not binding |
| HC11 | not binding (BLI) | | | not binding |
| HC21 | not binding (BLI) | | | not binding |

**Supp. Table 10. Model evaluation (nested CV); Folds for CV: k_inner = 5, k_outer=5**

| model_name | MSE | R2 | MSE | R2 | best_params_nonnested | most_selected_params_nested |
|---|---|---|---|---|---|---|
| | not nested | | nested | | not nested | nested |
| GP RBF | 0.0225 | 0.7605 | 0.0227 | 0.7589 | {'alpha': 1e-10, 'kernel': Matern(length_scale=1, nu=1.5), 'n_restarts_optimizer': 1} | {'alpha': 1e-10, 'kernel': Matern(length_scale=1, nu=1.5), 'n_restarts_optimizer': 1} |
| GP Matern | **0.0208** | **0.7816** | **0.0209** | **0.7804** | {'alpha': 1e-10, 'kernel': RBF(length_scale=1), 'n_restarts_optimizer': 1} | {'alpha': 1e-10, 'kernel': RBF(length_scale=1), 'n_restarts_optimizer': 1} |
| KernelRidge | 0.0494 | 0.5512 | 0.0511 | 0.5267 | {'alpha': 1.0, 'degree': 2, 'gamma': 0.1, 'kernel': 'polynomial'} | {'alpha': 0.1, 'degree': 2, 'gamma': 0.1, 'kernel': 'linear'} |
| RandomForest | 0.0292 | 0.7155 | 0.0302 | 0.7035 | {'fit_intercept': False} | {'fit_intercept': False} |
| LinearRegression | 2.73E+21 | 4.76E+22 | 1.47E+21 | -2.11E+22 | {'max_depth': 10, 'n_estimators': 200} | {'max_depth': 10, 'n_estimators': 200} |

**Supp. Table 11. Model evaluation (LOO CV)**

| Model | R2 | Corr_coef | MSE | params |
|---|---|---|---|---|
| GP RBF | 0.8412 | 0.9194 | 0.0216 | GaussianProcessRegressor(alpha=0.001, kernel=RBF(length_scale=1), random_state=1) |
| GP Matern | **0.8625** | **0.9313** | **0.0188** | GaussianProcessRegressor(alpha=0.001, kernel=Matern(length_scale=1, nu=1.5), random_state=1) |
| KernelRidge | 0.7360 | 0.8670 | 0.0360 | KernelRidge(alpha=0.1, degree=2, gamma=0.1, kernel='polynomial') |
| RandomForest | 0.8224 | 0.9211 | 0.0242 | RandomForestRegressor(max_depth=5, n_estimators=10, random_state=1) |
| LinearRegression | -4.93E+22 | -0.0736963 | 6.72E+21 | LinearRegression() |

**Supp. Table 12. Synthetic in silico designed antibody variants**

| Sample | $K_D$ [nM] | predicted $K_D$ [nM] | VH aa seq |
|---|---|---|---|
| tHC4.1 | 3.623 | 2.881 | QVQLQQSGAELVRPGASVTLSCKASGYTFTDYEMHWVKQTPVHGLEWIGAIDPETGSTAYNQNFKGKATLTADKSSSTAYMEFRSLPSEDSAVYYCTRDYYGSNYQAWFAYWGQGTLVTVSA |
| tHC1.1 | 1.294 | 0.854 | QVQLQQSGAELVRPGASVTLSCKASGYTFTDYEMHWVKQTPVHGLEWIGDIDPETGGTAYNQNFKGKATLTADKSSSTAYMEFRSLTSEDSAVYYCTRDYYGSNYQAWFAYWGQGTLVTVSA |
| tHC2.1 | 1.7655 | 1.364 | QVQLQQSGAELVRPGASVTLSCKASGYTFTDYEMHWVKQTPVHGLEWIGDIDPETGSTAYNQNFKGKATLTADKSSSTAYMEFRSLTSEDSAVYYCTRDYYGSNYQAWFAYWGQGTLVTVSA |
| tHC7.1 | 0.3307 | 3.378 | QVQLQQSGAELVRPGASVTLSCKASGYTFTDYEMHWVKQTPVHGLEWIGAIDPETGSTAYNQNFKGKATLTADKSPSTAYMEFRSLPSEDSAVYYCTRDYYGSNYQAWFAYWGQGTLVTVSA |
| tHC6.1 | 0.6152 | 1.205 | QVQLQQSGAELVRPGASVTLSCKASGYTFTDYEMHWVKQTPVHGLEWIGDIDPETGGTAYNQNFKGKATLTADKSPSTAYMEFRSLTSEDSAVYYCTRDYYGSNYQAWFAYWGQGTLVTVSA |
| tHC9.1 | 0.3507 | 0.334 | QVQLQQSGAELVRPGASVTLSCKASGYTFTDYEIHWVKQTPVHGLEWIGDIDPETGGTAYNQNFKGKATLTADKSSSTAYMEFRSLTSEDSAVYYCTRHYYGSNYLAWFAYWGQGTLVTVSA |
| tHC8.1 | 0.4885 | 0.396 | QVQLQQSGAELVRPGASVTLSCKASGYTFTDYEMHWVKQTPVHGLEWIGDIDPETGGTAYNQNFKGKATLTADKSSSTAYMEFRSLTSEDSAVYYCTRHYYGSNYLAWFAYWGQGTLVTVSA |
| tHC10.1 | 0.4007 | 0.327 | QVQLQQSGAELVRPGASVTLSCKASGYTFTDYEMHWVKQTPVHGLEWIGDIDPETGGTAYNQNFKGKATLTADKSSSTAYTEFRSLTSEDSAVYYCTRHYYGSNYLAWFAYWGQGTLVTVSA |

**Supp. Table 13: Model evaluation on the novel variants**

| Model | R2 | Corr_coeff | MSE | Model_params |
|---|---|---|---|---|
| GP RBF | -0.2327 | 0.4936 | 0.2140 | GaussianProcessRegressor(kernel=RBF(length_scale=1), random_state=1) |
| GP Matern | **-0.1686** | 0.4741 | **0.2029** | GaussianProcessRegressor(kernel=Matern(length_scale=1, nu=1.5), random_state=1) |
| KernelRidge | -0.5551 | 0.3912 | 0.2700 | KernelRidge(alpha=0.1, degree=2, gamma=0.1, kernel='polynomial') |
| RandomForest | -0.4999 | **0.5465** | 0.2604 | RandomForestRegressor(max_depth=5, n_estimators=10, random_state=1) |

**Supp. Table 14. Model evaluation (LOOCV) with training dataset + novel variants;**

| Model | R2 | Corr_coeff | MSE | Model_params |
|---|---|---|---|---|
| GP RBF | 0.3430 | 0.6139 | 0.0942 | GaussianProcessRegressor(alpha=0.01, kernel=RBF(length_scale=1), random_state=1) |
| GP Matern | 0.3507 | 0.6255 | 0.0931 | GaussianProcessRegressor(alpha=0.01, kernel=Matern(length_scale=10, nu=1.5), random_state=1) |
| KernelRidge | 0.2049 | 0.6075 | 0.1140 | KernelRidge(alpha=1.0, gamma=0.1, kernel='polynomial') |
| RandomForest | **0.3791** | **0.6314** | **0.0890** | RandomForestRegressor(max_depth=5, n_estimators=200, random_state=1) |

**Supp. Table 15. Model evaluation on the novel variants (outlier removed)**

| Model | R2 | Corr_coeff | MSE | Model_params |
|---|---|---|---|---|
| GP RBF | **0.8669** | 0.9378 | **0.0236** | GaussianProcessRegressor(alpha=0.01, kernel=RBF(length_scale=1), random_state=1) |
| GP Matern | 0.8348 | 0.9238 | 0.0293 | GaussianProcessRegressor(alpha=0.01, kernel=Matern(length_scale=10, nu=1.5), random_state=1) |
| KernelRidge | 0.7126 | 0.9075 | 0.0509 | KernelRidge(alpha=1.0, gamma=0.1, kernel='polynomial') |
| RandomForest | 0.8040 | **0.9398** | 0.0347 | RandomForestRegressor(max_depth=5, n_estimators=200, random_state=1) |
